# Molecular Polymorphism of tau aggregates in Pick’s disease

**DOI:** 10.1101/2024.11.19.624253

**Authors:** Jiliang Liu, Theresa Connors Stewart, Bradley T. Hyman, Manfred Burghammer, Marine Cotte, Lee Makowski

## Abstract

Tau protein plays a central role in many neuropathies. The trajectory by which tau spreads through neural networks is disease-specific but the events driving progression are unknown. This is due in part to the challenge of characterizing tau aggregates *in situ*. We address that challenge using *in situ* micro-x-ray diffraction (µXRD) and micro-X-ray fluorescence (μXRF) to examine tau lesions in the brain of a 79-year-old male with dementia. Neuropathological examination revealed classical forms of tau in the hippocampal formation: extensive Pick bodies in the granular layer; modest numbers of neurofibrillary tangles and dystrophic neurites in the CA4 and hilus. µXRD indicated that Pick bodies are low in fibril content, whereas neurofibrillary lesions within adjacent tissue exhibit far greater density of fibrillar tau. μXRF demonstrated elevated levels of zinc, calcium and phosphorous in all tau-containing lesions whereas sulfur deposition was greatest in lesions exhibiting high fibrillar content. Correlation of lesion morphology with anatomical localization, tau fibrillation and differential elemental accumulation suggests tau fibrils generate biochemically distinct microenvironments that influence lesion morphology, tau seed formation and spreading.

## Introduction

Pick’s disease, also known as frontotemporal dementia (FTD) or frontotemporal lobar degeneration with tau pathology (FTLD-tau), is a progressive, neurodegenerative disorder that impacts the brain’s frontal and temporal lobes. Aggregates composed of tau protein are closely associated with the progression of dementia in Pick’s disease. Pick bodies, intracellular spherical inclusions unique to Pick’s disease, build up inside neurons that take on a swollen, balloon-like shape. Within these lesions, fibrillar forms of tau contain only 3R isoforms and exhibit a unique cross-sectional core structure.(1) In about 10% of Pick’s disease cases, including the one we report on here, there is additional Alzheimer’s disease pathology, indicated by the presence of neurofibrillary tangles (NFTs).(2) The presence of Pick bodies and NFTs within a single tissue section provided the opportunity to compare the properties of two different forms of tau inclusion using an advanced set of µXRD and µXRF tools.

*In situ*, in addition to the well studied cross-β fibrils, tau aggregates may exhibit relatively diverse morphologies including oligomeric forms, amorphous or ring-like aggregates, small oligomers, or monomeric species.(3) Most studies of the molecular structure of these aggregates utilize purified tau and cannot provide positional information on structural variation within tissue. But the spatial distribution of fibrillar and smaller aggregates of tau in tissue may well influence the processes underlying seeding and spreading of tau.(3) Therefore, generating information on the positional variation of aggregate structure is a high priority. Here, we use scanning micro X-ray diffraction (µXRD) to study the structural variations of tau within tissue and micro X-ray fluorescence (μXRF) microscopy to assess the accumulation of specific elements within the lesions. In this approach, a histological tissue section is raster scanned in front of a micro-focused X-ray beam and XRD and XRF data are collected simultaneously to characterize position-specific variation of molecular structure and metal accumulation with subcellular resolution. Immunohistochemistry of serial sections is used to identify the major constituents of lesions.

The size and shape of pathological lesions in human brain tissue can be determined by mapping the intensity of wide-angle X-ray scattering across a thin section. (4) Higher intensity indicates denser packing of macromolecular constituents and most pathological lesions appear to have higher packing density than the surrounding proximal tissue. The size and shape of features identified in this way correspond closely with those of tau-staining lesions observed by immunohistochemistry in the associated serial sections, providing strong evidence of their identity as tau-rich lesions. Other structural features of tissue, such as vascular walls also have greater density than the surrounding tissue and these can be identified by their characteristic shapes in maps of scattering intensity.

All neuropathological fibrils have a distinctive cross-β fibrillar structure which gives rise to a stereotypic X-ray scattering ‘fingerprint’. In a cross-β structure, β-strands of protein extend across the fibril, perpendicular to the fibril axis, forming β-sheets of indeterminant length. The structure acts as a diffraction grating, generating strong X-ray scattering at angles that correspond to the axial distance between β-strands (4.7 Å) and the transverse distance between the β-sheets (10 Å).(5,6) X-ray scattering from a tissue sample measured with conventional diffractometers will not produce high quality signal from fibrils embedded in the tissue because the scattering volume is so large as to encompass heterogeneous tissue structures beyond the boundaries of the lesion. This results in scattering patterns that are the sum of scattering from a heterogeneous mixture of structures that obscures the scattering from lesions and precludes interpretation on the basis of individual molecular structures. The use of a microbeam from a synchrotron source allows collection of a scattering signal from a volume sufficiently small that it falls entirely within a lesion and is dominated by a single constituent, thereby giving rise to scattering patterns that can be interpreted in terms of the structure of that constituent.(7,8,9) This makes possible study of the molecular structure of tau aggregates such as neurofibrillary tangles (NFTs), Pick bodies, and dystrophic neurites that vary in size from a few microns to tens of microns. Correlation with immunohistochemistry of serial sections makes it possible to positively identify the principal constituents of these lesions and to place the structural attributes of these aggregates in the broader context of the cellular organization of the tissue.

As will be demonstrated here, data in the wide-angle portion of the X-ray patterns (WAXS) can be used to assess the degree of fibrillation of tau, while intensity in the small angle X-ray scattering regime (SAXS) can provide insights into the structure and packing of tau fibrils. Simultaneous collection of X-ray fluorescence (XRF) data makes possible correlation of the deposition of selected metal and non-metal elements with the different levels of tau accumulation and fibrillation revealed by the scattering data. When combined with correlated immunohistochemistry of serial sections, these data provide a basis for associating distinct tau structures with different cell types distributed within specific regions of the brain. The structural information derived directly from tissue can provide insights in the process of seeding and spreading of tau in Pick’s disease to supplement that derived through studies of isolated materials.

## Results

### Simultaneous collection of μXRD and μXRF maps

**Figure 1a** diagrams the experimental arrangement used to simultaneously collect X-ray scattering and X-ray fluorescence data across a tissue section raster scanned in front of a micro-focused X-ray beam. Details about the experimental set-up, sample preparation and data processing are given below in the methods section. In brief, thin sections of human brain tissue from a Pick’s disease subject were prepared as previously described (4) and spread on 1 μm thick Si3N4 membranes. These sections were scanned at the micro-end-station of the beamline ID13, at the European Synchrotron Radiation Facility (ESRF) with an X-ray beam having full width half maximum (fwhm) of 2.5 μm. 2D regions of interest (ROI) ∼1500 µm on a side were defined for each section and XRD and XRF data were collected on a square, 2.5 µm grid across these ROIs (roughly 600×600 frames constituting 360,000 diffraction patterns). X-ray patterns from tissue sections are usually circularly symmetric, or nearly so. Consequently, we circularly averaged the patterns to improve signal-to-noise ratio. As shown in **Figure 1b**, the circularly averaged data includes strong intensity in the small-angle (SAXS) regime and distinctive broad peaks at scattering angles that correspond to periodicities of 4.7 Å and 10 Å. Fixed tissue from many organs will exhibit broad peaks at these spacings due to residual secondary structure in partially denatured proteins. Scattering from regions rich in fibrillar material (blue curve in **Figure 1b**) gives rise to a relatively sharp, pronounced reflection at 4.7 Å spacing located on a broad scattering peak arising from tissue constituents interspersed with the fibrils. Lesions with low fibrillar content produce a relatively weak 4.7 Å peak (orange curve). Tissue regions devoid of tau fibrils or aggregates generate a broad wide-angle peak (green curve) that is usually weaker than that observed from tau-containing lesions. Analysis of the scattering patterns in terms of the structure of the molecular constituents of the scattering volume is described in **Figure S1**. Additional experimental and computational details are given in the methods section. Simultaneously collected XRF signal identified increased fluorescence intensities of calcium and zinc associated with lesions containing tau as seen in **Figure 1c**.

**Figure 1.**
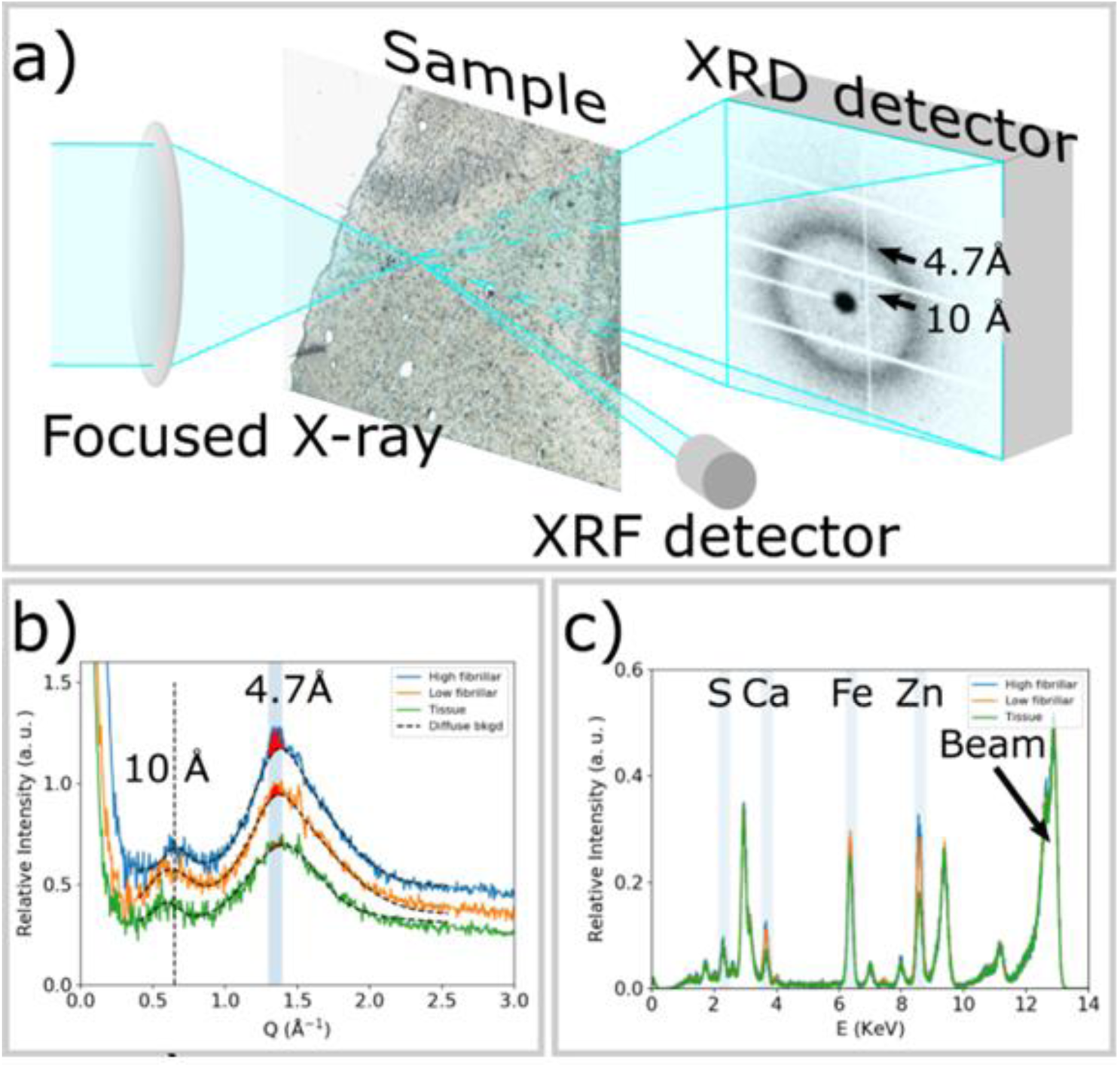
**a)** Experimental scheme for collecting µXRD and µXRF data simultaneously. The X-ray beam is focused to 2.5 μm by the lens and the sample is raster scanned with step size of 2.5 μm. µXRD from tau lesions gives rise to a prototypical cross-β fiber diffraction pattern, which is dominated by reflections at scattering angles corresponding to periodicities of 10 Å and 4.7 Å. An XRF detector is placed nearly perpendicular to the sample to simultaneously collect XRF signal. **b)** Azimuthally averaged scattering patterns from a tau lesion with significant high-fibrillar tau (blue); a tau-containing lesion containing low-fibrillar tau (orange) and a region of tissue exhibiting no tau pathology (green). High-fibrillar tau gives rise to a pronounced 4.7 Å peak, the intensity of which can be estimated by subtracting a smooth background and integrating the remaining intensity (red fill). Short fibrils, or tau aggregates that are low-fibrillar may give rise to a weak peak at 4.7 Å spacing that can also be estimated through subtraction of a smooth background. Surrounding tissue also gives rise to broad scattering peaks at ∼ 10 Å and 4.7 Å spacing, but lacks the additional pronounced features at 4.7 Å. **c)** μXRF spectra from a tau lesion with significant high-fibrillar tau (blue); a tau-containing lesion with little or low-fibrillar tau (orange) and a region of tissue exhibiting no tau pathology (green). The spectral line exhibits patterns of chemical elements, including sulfur (S) with Kα edge at 2.31 KeV, calcium (Ca) at 3.64 KeV, iron (Fe) at 6.4 KeV and zinc (Zn) at 8.64 KeV. The peak at 13 KeV arises from Rayleigh and Compton scattering of the incident X-ray beam. The maps of elemental distributions are calculated by integrating the spectrum over a range of ± 0.1KeV around the corresponding Kα energy for each pixel. Intensity of peaks in the XRF spectrum reveal a relative degree of deposition of each element within the tissue sample.

### Distribution of tau aggregates in tissue

Although tau is present throughout the cortex in Pick’s disease, the dentate gyrus (DG) of the hippocampus frequently contains a high density of tau lesions.(10) The DG granule cell layer typically is spared in Alzheimer disease, but in Pick’s disease it may contain a number of different types of tau deposits. These include Pick bodies, exhibiting a prototypic round morphology, often heavily deposited in the granular layer; and thin filaments of tau heavily deposited within dystrophic neurites in the hilus. In about 10% of Pick’s disease cases, including the one we report on here, there are features of concurrent Alzheimer’s disease pathology that has fibrillar neurofibrillary tangles (NFTs), found inside neuronal and pyramidal cells (including those in the hilus, adjacent to the granule cells of the DG).(11) **Figure 2** compares images of the tissue produced by different imaging modalities. **Figure 2a** is an optical micrograph of a portion of the dentate gyrus immunostained for Aβ, but showing little evidence for the presence of Aβ lesions. **Figure 2b** is the corresponding serial section immunostained for tau that reveals an abundance of Pick bodies in the granular layer. These are identified by their dark brown staining and round shape as highlighted in the upper inset. The hilus and Cornu Ammonis region 4 (CA4) are composed largely of pyramidal cells.(12,13) Within the hilus region, tau deposits are largely dystrophic neurites and NFTs that exhibit a characteristic elongated teardrop morphology as seen in the lower inset. **Figure S2a** provides a wider field of view of the serial section immunostained for tau showing two dense granular layers separated by the hilus region that connects to the CA4 and CA3 regions of the hippocampus. These granular layers exhibit particularly dense deposition of Pick bodies.

**Figure 2.**
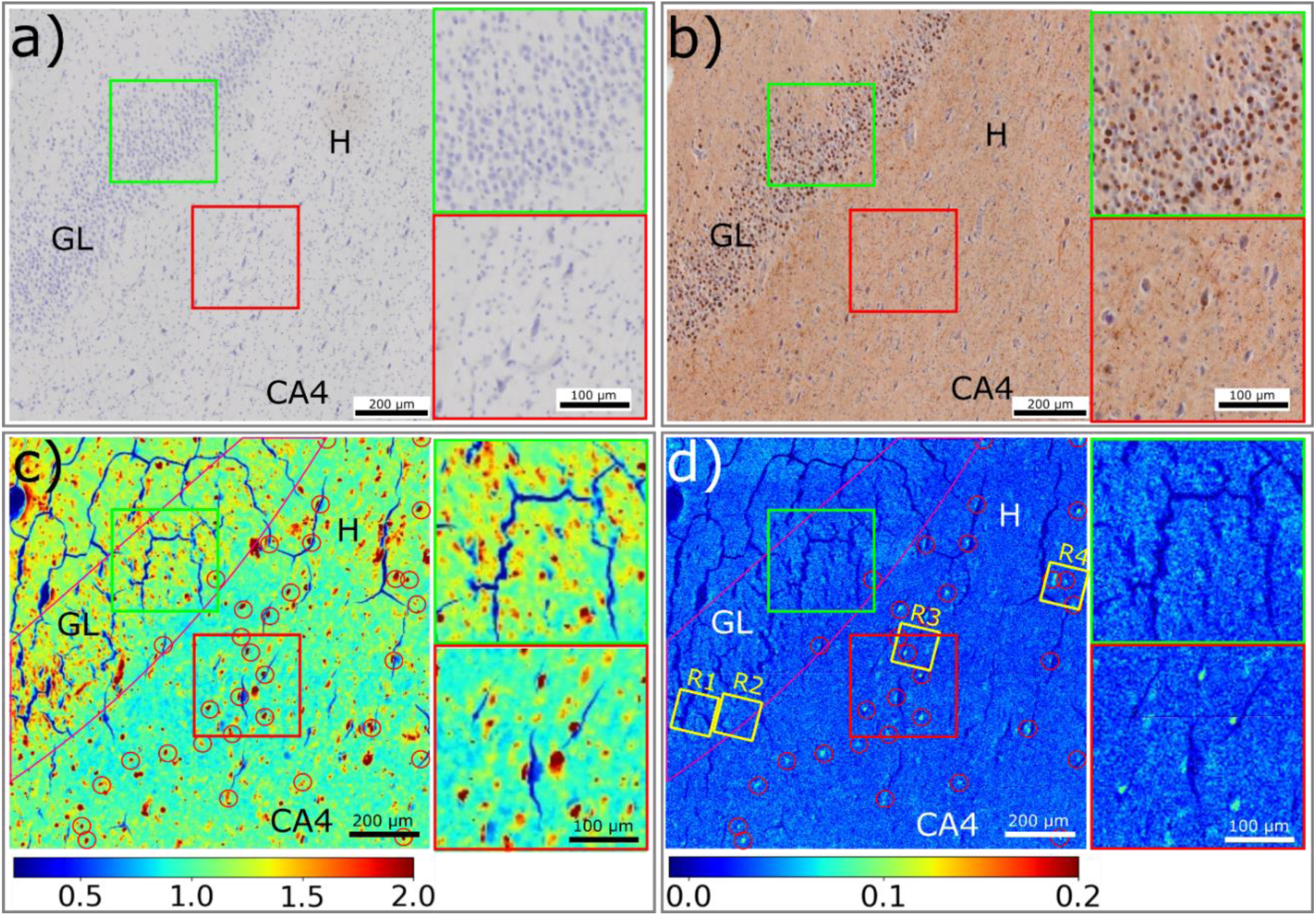
µXRD analysis of a region of a Dentate Gyrus including the Granular Layer (GL), Hilus (H) and the Cornu Ammonis region 4 (CA4) of the hippocampus. **a)** Image of a serial section stained for Aβ corresponding to the region of the unstained section scanned by XRD. Different cell morphologies, including granule cell and pyramidal cell, are present. **b)** Image of a serial section stained for tau. The granular layer is dominated by Pick bodies, while tau lesions from dystrophic neurites and pyramidal cells are distributed across the hilus region. Two inserts show the distribution of tau lesions in granular layer (green box) and hilus region (red box). **c)** The distribution of macromolecular density is calculated by integrating the small-angle X-ray scattering intensity within the range of 0.05 – 0.25 Å^-1^ of Q. The distribution of macromolecular density exhibits dense features in both the granular layer, CA4 and hilus and includes features that are low in fibrillar content and those that exhibit a high concentration of fibrillar tau (indicated by red circles). **d)** The distribution of fibrillar tau is determined by the integral intensity of patterns after diffuse background subtraction. Regions containing oligomeric or low fibrillar aggregates appear as light blue; high fibrillar lesions as bright blue (marked by red circles). Two inserts on the right show the distribution of aggregates in the Granular layer (green box) and fibrils in the Hilus region (red box). **R1**-**R4** are four ROIs in which high resolution XRF images were collected as detailed in Figure 5. The relative intensity of color scale for c) and d) is given in arbitrary units.

### The aggregation state of tau is specific to cellular environment

The location of tau lesions, be they Pick bodies, neurofibrillary tangles or dystrophic neurites can be distinguished by the intensity distribution of X-ray scattering in the wide-angle regime. As shown in **Figure 1b**, this scattering can be divided into two classes – broad, diffuse scattering which all proteinaceous macromolecular structures contribute to; and pronounced 4.7 Å scattering which is characteristic of cross-β fibrillar structures such as those formed by tau. The broad wide-angle scattering is observed in all fixed tissue but is usually observed to be more intense in Aβ or tau lesions.(4,14) This indicates that pathological lesions are more densely packed with macromolecules than the surrounding tissue. **Figure 2c** shows the distribution of macromolecular density across the ROI as estimated from the intensity of scattering in the range 0.05 < Q < 0.25 Å^-1^. The size, shape and density of the features apparent in this mapping correlate well with the corresponding features of Pick bodies observed by immunohistochemistry of serial sections as seen in **Figure 2b**. While immunohistochemistry confirms the presence of tau, the relative abundance of tau and other macromolecules in these lesions remains unknown.

The location of cross-β fibrillar structure can be determined by estimating the intensity of the pronounced 4.7 Å peak (as detailed in **Figure S1**) resulting in a map as shown in **Figure 2d**. By comparing the dense features in **Figure 2c** with the positions of fibrillar lesions (indicated by red circles) in **Figure 2d**, the positions of tau lesions with greater or lesser fibrillar content can be determined. In the CA4 and hilus, fibrillar tau is associated with many but not all lesions. In the granular layer, most lesions exhibit little or no fibrillar structure. While electron microscopy (and cryo-electron microscopy of isolated tau filaments) have demonstrated the presence of fibrillar tau in Pick bodies (15), our observations indicate that the content of fibrillar tau in Pick bodies in the granular layer is relatively low and that most of the tau present must be in disordered or amorphous aggregates or fibrillar fragments too small to be detected by X-ray scattering. By contrast, fibrillar structures within typical AD related neurofibrillary tangles present in pyramidal cells in the same tissue, are readily detected, reflecting a more robust fibrillar phenotype.

To summarize, the distribution of small angle scattering intensity from the granular layer, molecular layer and hilus (**Figure 2c**) indicates a distribution of tau that closely corresponds to that identified by immunostaining of the serial section in **Figure 2b**. However, the distribution of high ***fibrillar tau*** (**Figure 2d**) as assessed from the intensity of the 4.7 Å peak, is distinct from the distribution of total tau with the degree of fibrillation in the Pick body containing granular layer significantly less than in the tangle containing hilus region. Additional analysis of the 4.7 Å peak provides clearer visualization of the fibrillar distribution as shown in **Figure S3**.

### Fibrillar deposits of tau exhibit a hierarchical organization

In addition to the 4.7 Å peak, as shown in **Figure 3a**, SAXS intensities from fibrillar lesions exhibit features not seen in scattering from tissue or regions of low fibrillar content. Both low fibrillar and high fibrillar tau give rise to stronger SAXS intensity than does tissue, as shown in the mapping of SAXS intensity in **Figure 2c**. However, the linear slope of the log-log scaled SAXS profile of low fibrillar tau indicates that the tau and macromolecular aggregates are highly polydisperse in size. In contrast, scattering from regions with a high concentration of fibrils exhibit distinctive peaks or shoulders in the SAXS regime (**Figure 3a**). This modulation of SAXS intensity is only observed in patterns that also exhibit pronounced reflections at 4.7 Å spacing, confirming they are due to the presence of cross-β fibrils. These peaks provide information about the transverse (cross-sectional) morphology of the fibrils and their higher-order spatial organization. Quantitative analysis of these fibril-specific features was carried out after subtraction of background from the amorphous tissue and/or voids that are interspersed with fibrils in the scattering volume.(17).

**Figure 3.**
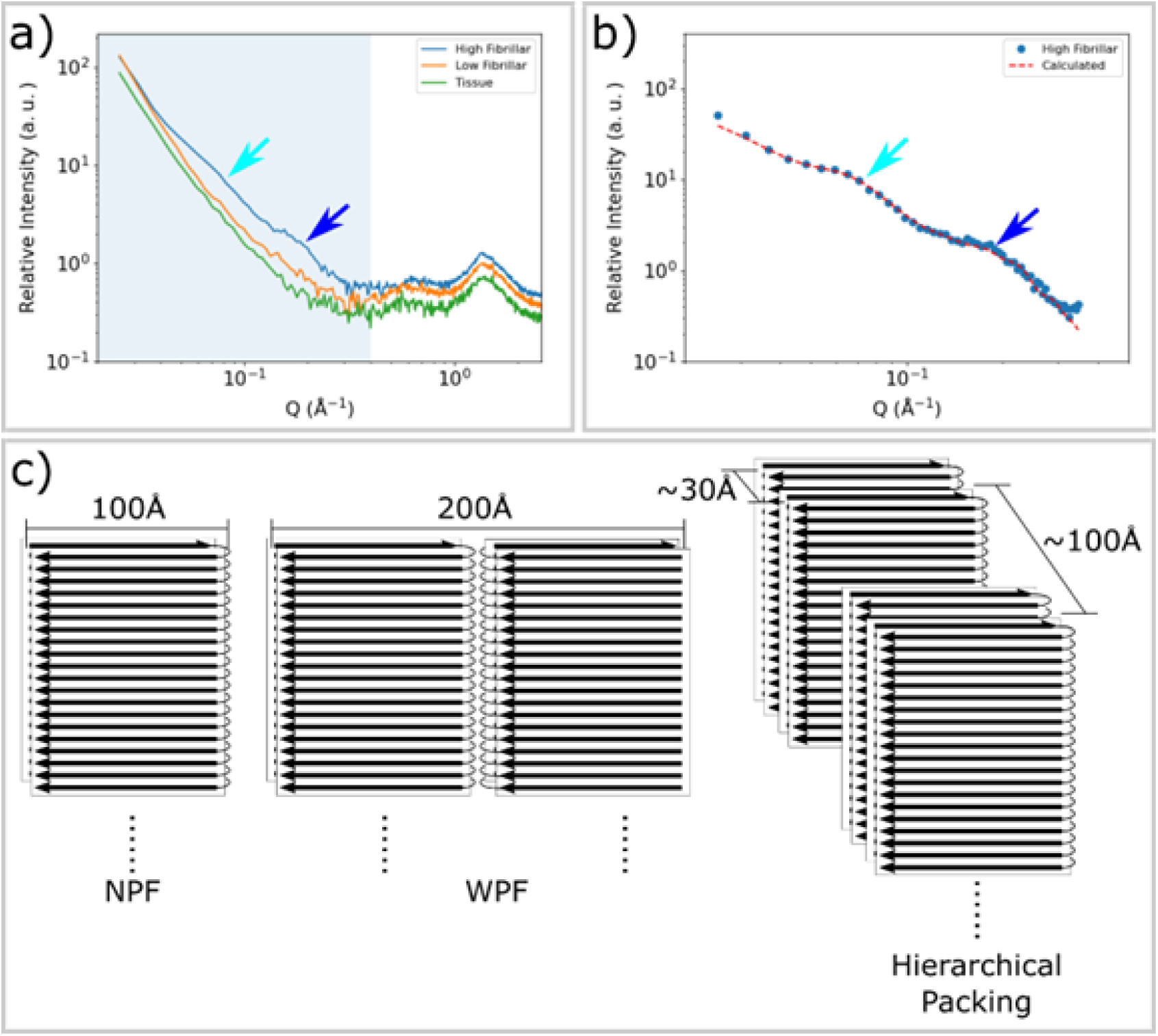
SAXS from fibrillar tau exhibits distinctive peaks. **a)** The SAXS profiles from tissue devoid of tau aggregates (green), containing low fibril content (orange) and having high levels of fibrillar tau (blue) as identified by the intensity of the pronounced WAXS reflection at 4.7 Å spacing. Patterns exhibiting strong cross-β related features in the WAXS regime (Q ∼ 1.36 Å^-1^) also exhibit distinctive features in the SAXS regime, highlighted by cyan and blue arrows at Q ∼ 0.07 Å^-1^ and ∼ 0.2 Å^-1^. **b)** Background-subtracted SAXS intensities correspond well with those calculated from a hierarchical model of fibrillar organization exhibiting hierarchical packing with limited variation in fibril-fibril distances. **c)** A diagram of the narrow pick filament (NPF), the wide pick filament (WPF) and the model of hierarchical packing used to fit the observed data. Inter-fibrillar distances of 30 Å and 100 Å are representative of the polymorphic hierarchical organization indicated by the shape of the SAXS scattering as detailed in **Figure S4**.

Simplified structural models for the tau fibril, **Figure 3c**, were constructed on the basis of the structure of tau filament cores as determined by cryoEM studies of material extracted from Pick’s disease subjects.(1) However, models based on the structure of individual fibrils (either narrow Pick filament (NPF) or wide Pick filament (WPF)) failed to predict the observed intensities. A simplified cylinder model (18) also failed to reproduce details of the observed experimental intensity. As detailed in the Supplementary Material (**Figure S4**), in order to successfully reproduce the observed intensities, a polymorphic hierarchical organization of fibrils was postulated with characteristic fibril-fibril distances of 30 Å (+/- 15%) and 100 Å (+/- 30%). This model resulted in predictions of scattered intensity with characteristics that closely correspond to those observed (**Figure 3b**). The relatively large fibril-fibril distances characteristic of this packing appears to be required in order to accommodate the intrinsically disordered ‘fuzzy coat’ of tau (e.g.(2)) that accounts for about 75% of the mass of the tau molecule in neurofibrillary tangles.

### Variation of Elemental Accumulation with Aggregation state of Tau

Simultaneous collection of μXRF and µXRD data allows correlation of elemental distributions with tau accumulation and fibrillation. The 2.5 μm μXRF images for Zn and Ca have signal-to-noise ratios sufficient for mapping the localization of these elements across the ROI scanned by μXRD (**Figure 4**). Both zinc and calcium (**Figure S7**) accumulate in essentially all tau lesions whether tau fibrils are present or not. Calcium and zinc have also been found to co-locate with Aβ within plaques of the brains from AD subjects.(19) To get better detection of low-Z elements, additional µXRF maps were acquired at the ID21 beamline, which operates under vacuum to make possible the use of low-energy X-rays (2.1-11keV) required to detect these elements (see details below in the methods section) (**Figure 5** and **Figure S9**). **Figure 5** is a comparison of the distribution of Zn, P, Ca, and S (as determined by μXRF) with the presence of tau aggregates and fibrils in four regions as designated in **Figure 2d**. In this Figure, **regions 1 and 2** are in the granular layer, where tau is low in fibrillar content and **regions 3 and 4** are in the hilus where lesions are dominated by fibrillar tau. Like Zn and Ca, P appears to co-locate with tau independent of fibrillization state, suggesting that tau phosphorylation is not detectably different between fibrillar and non-fibrillar forms of tau in this tissue context, in accord with mass spectroscopy data that show that tau is hyperphosphorylated (although with different patterns) in both these lesions.

**Figure 4.**
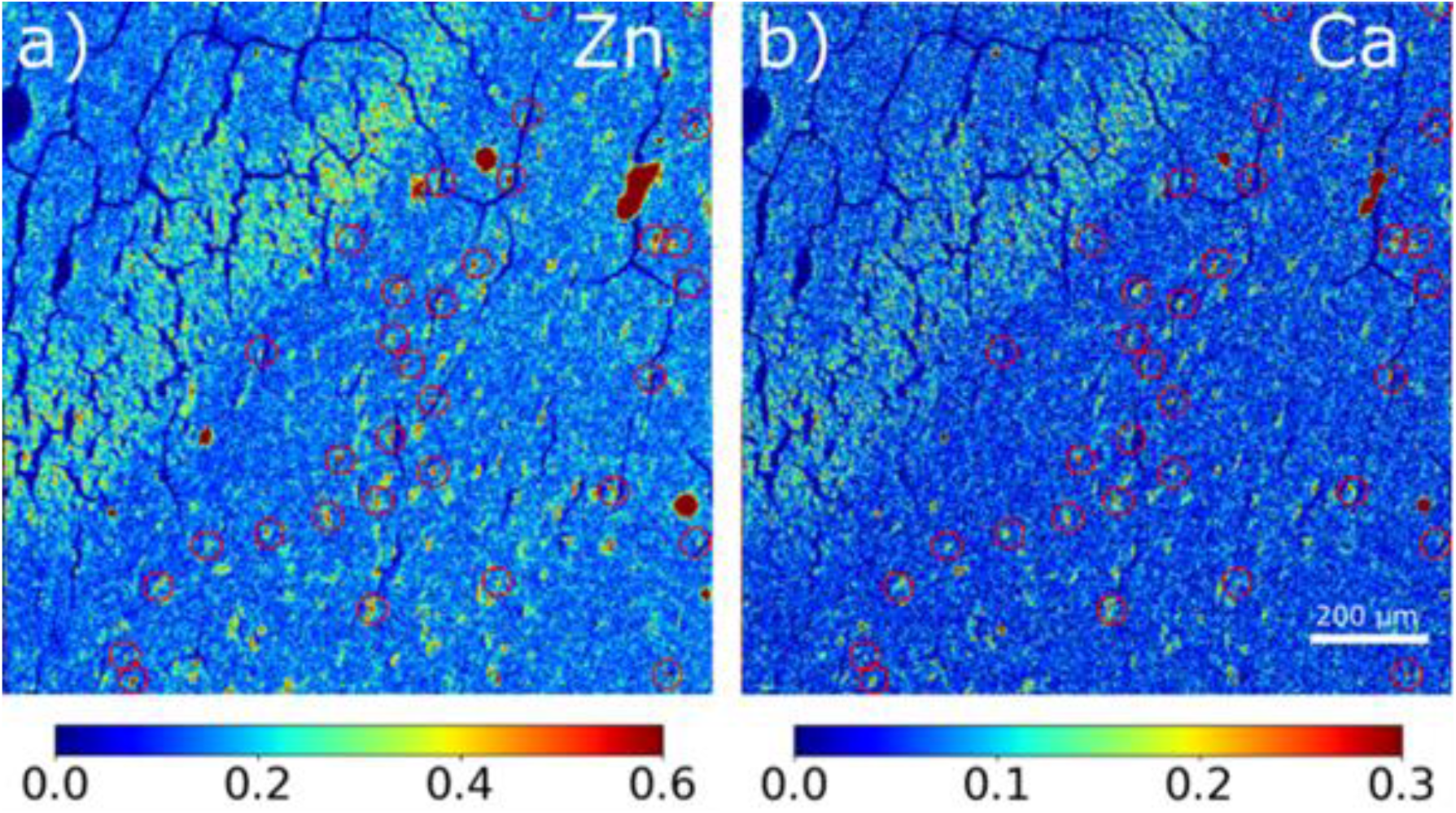
Distribution of X-ray fluorescence signal from **(a)** zinc and **(b)** calcium. Red circles indicate the locations of fibrils as determined from µXRD. The relative intensity of color scale bars is in arbitrary units.

**Figure 5.**
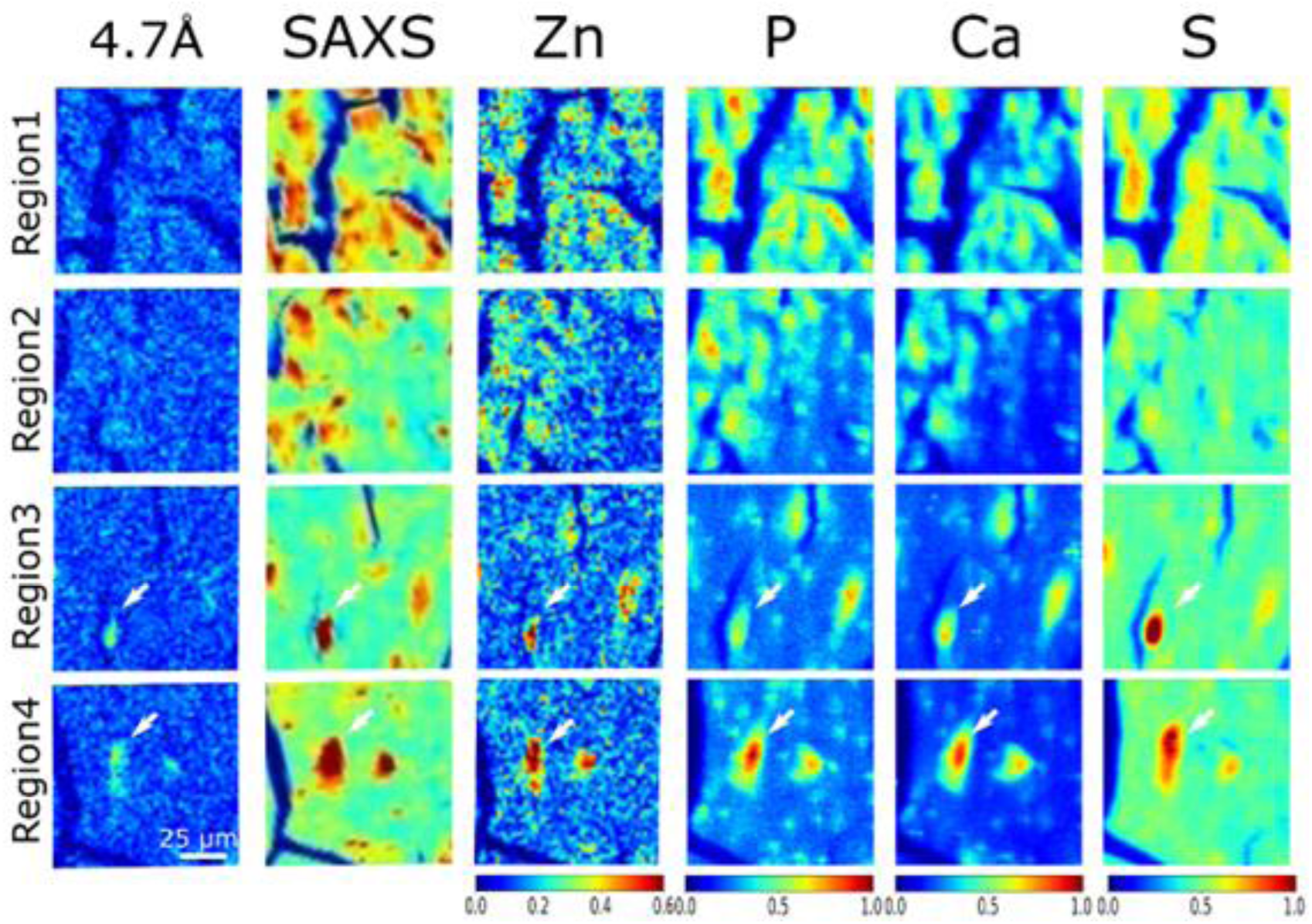
Correlation of the distribution of tau fibrils (left column), integral intensity of SAXS from macromolecular deposition (second column from left) and distribution of select elements. **Regions 1-4** correspond to those highlighted as yellow boxes in Figure 2d and **Extended Data Figure S6** and **S8**. **Regions 1 and 2** are in the granular layer, where tau is low in fibrillar content as indicated by µXRD. **Regions 3 and 4** are in the hilus where lesions are dominated by fibrillar tau. The white arrows in **Regions 3 and 4** highlight the fibrillar tau identified by the 4.7 Å, β-strand reflection. The relative intensity of color scale bars is given in arbitrary units.

In contrast, the deposition of sulfur appears to be greater in lesions containing fibrillar tau than in lesions in the granular layer that have relatively lower fibrillar content. Sulfated glycosaminoglycans are known to induce the formation of tau filaments *in vitro*.(20,21) Observation of a correlation of tau fibrillation with high sulfur deposition *in situ* provides support for the hypothesis that, consistent with *in vitro* observations, glycosaminoglycans may actively promote tau fibrillation in tissue.(22,23)

## Discussion

A combination of immunohistochemistry, scanning µXRD, and μXRF has been used to assess the distribution of tau aggregate types across a single section of human brain tissue, revealing that the level of tau fibrillization within a lesion correlates with its gross morphological characteristics. Pick bodies in the granular layer of the DG contain relatively low concentrations of fibrils, whereas NFTs in the adjacent CA4 and hilus regions have a much higher fibrillar content. The presence of tau fibrils in Pick bodies was demonstrated decades ago by electron microscopy.(15) However, our XRD studies indicate that the density of fibrils within Pick bodies is far lower than in NFTs in adjacent brain regions. Within NFTs, fibrils are packed so tightly that they form domains with a relatively well-ordered hierarchical organization. This dense packing may influence the accumulation or exclusion of other macromolecular species or elements, thereby affecting the microenvironment within the lesions.

We have used XRF microscopy to show that some elements accumulate in tau-containing lesions at higher concentrations than in the surrounding tissue. Tissue fixation and preparation may wash out elements not bound to immobilized species, suggesting that most detected elements are specifically bound to macromolecular constituents. Levels of Zn, Ca and P appeared higher in lesions than in the surrounding tissue, with no overt preference for lesion type. In contrast, no accumulation of Na, Mg, and Al was observed in any tau-containing lesion.

Accumulation of phosphorus in tau-containing lesions was expected due to the role that hyper-phosphorylation of tau appears to have in disease progression. μXRF imaging confirms the accumulation of phosphorus in essentially all tau lesions examined with no apparent preference for lesions with higher or lower levels of tau fibrillation. This suggests that phosphorylation of tau may play a passive role in tau fibril assembly. In contrast, sulfur appears to accumulate to higher levels in fibrillar lesions than in lesions with low levels of fibrillar tau, consistent with the hypothesis that sulfur-containing compounds may play a role in tau fibrilization (20).

The correlation between tau fibrillation levels, elemental deposition, and gross lesion morphology suggests that the biochemical milieu within tau lesions may be influenced not only by the accumulation of tau but also by the degree of fibrillar tau formation. Similar to how biological condensates create local microenvironments that promote distinct biochemical processes (24), high concentrations of tau may alter the overall composition of lesions. This is particularly evident for elements traceable by μXRF, as demonstrated in our study. While it is unsurprising that the constituents of a lesion differ from the surrounding tissue, the differential behavior of lesions containing fibrillar versus non-fibrillar tau aggregates might be considered unexpected.

While cryo-electron microscopy approaches clearly show the fibrillar nature of material from both Pick bodies and neurofibrillary tangles with atomic resolution, they are not able to provide anatomical context or information about elemental composition in the same way as correlated µXRD and μXRF. These data demonstrate that Pick bodies appear to be substantially less fibrillar than neurofibrillary tangles, which is in accord with the observation that Pick bodies are less prominently stained with β-pleated sheet dyes like Thioflavin S or Congo Red. Moreover, the current data highlight differential association of several elements - both positively and negatively charged - with either Pick bodies or neurofibrillary lesions, with implications for their contribution to tau aggregation in different cellular milieus. The ability to map specific aggregation states of tau, as demonstrated here, offers a new approach to studying tau distribution and structural polymorphism.

## Materials and Methods

### Sample preparation

Brain tissue was prepared at the Massachusetts Alzheimer’s Disease Research Center. Tissue was formalin fixed using a standard neuropathological processes and sectioned to a thickness of 20 μm as previously described.(4) The brains collected by MADRC are handled, dissected, and stored in a uniform fashion. Brains are divided in half with one half fixed, processed in paraffin for neuropathological analysis and immunohistochemical studies and the other half generally snap-frozen in coronal slices. Regions of interest are cut from the fixed hemisphere, blocked into cassettes and stored in 10% formalin. Once selected for examination, a block is soaked in 98% formic acid for one hour. A tissue processor is then used to wash the tissue with formalin, and then a series of ethanol and then xylene solutions overnight. The tissue is then embedded in paraffin and cooled to harden the wax. Tissue is held at 5°C for sectioning into 20 μm thick sections. Three serial sections are collected, including one that is left unstained for x-ray analysis, one for immunostaining for tau and one for Aβ.

Unstained sections were prepared thicker than conventional histological sections in order to increase the volume of material irradiated thereby improving the signal-to-noise ratio of the scattering patterns collected. Tissue sections were mounted on 5 x 5 mm^2^ 1 μm thick Si3N4 membranes and affixed to sample holders customized for the ID13 beam line at ESRF.

### Simultaneous collection of µXRD and µXRF maps

Scanning µXRD was conducted at beam line ID13, at the European Synchrotron Radiation Facility (ESRF). The monochromatic beam was focused to a Gaussian profile with fwhm of 2.5 µm in both the horizontal and vertical directions, using compound refractive lenses. To reduce secondary radiation damage, a fly scanning mode was applied with scan step of 2.5 µm across a square grid. The energy of X-ray beam was 13 KeV, with 50 ms exposure time and 200 mA storage ring current. The microdiffraction patterns were collected in transmission mode by an Eiger 4M detector.

XRF spectra from elements heavier than Ca including Zn were collected with a Vortex EM detector installed almost normal to the path of the X-ray beam and pointing in the direction of the sample (**Figure 1**). Data processing was conducted using a custom program developed for scanning µXRD. XRF spectra have been analyzed using the PyMca program.(25)

### Mapping distribution of tau aggregates

The distribution of tau lesions within a fixed thin section was mapped using the intensity of scattering integrated over the range (0.05 Å^-1^ < Q < 0.25 Å^-1^). Intensity in this regime is approximately proportional to the electron density contrast of the macromolecular material within the scattering volume. When scanning a section of uniform thickness this is associated with the density of material in the tissue. There is substantial evidence that the density of macromolecular scattering material is greater in lesions (Aβ or tau) than in the surrounding material.(4,14) Other anatomical features such as vascular walls also have greater density than the surrounding tissue, and these are usually readily identified from their size and shape. Consequently, a map of scattering intensity across an ROI will allow the distribution of pathological lesions to be derived. The intensity in the range (0.05 Å^-1^ < Q < 0.25 Å^-1^) was used to calculate the map in **Figure 2c** because the intensities and signal-to-noise ratio are greater than at larger Q. Scattering intensities at smaller angles are impacted by scattering from voids formed during tissue processing and dehydration (17) and cannot be used for this purpose.

### Mapping the distribution of tau fibrils

Cross-β structure gives rise to a pronounced peak at ∼ 4.7 Å spacing (Q = 1.34 Å^-1^). As diagrammed in **Figure 1b**, the relative abundance of fibrillar tau was estimated by subtracting the broad diffuse background scattering and integrating the difference intensity over the Q range - 1.32 Å^-1^ to 1.4 Å^-1^. The smooth background curves (dashed lines) are calculated as a sum of Pseudo-Voigt functions and represent an estimate of the scattering of tissue, which is comprised largely of fixed, partially denatured macromolecules.

In X-ray scattering, the width of a peak scales inversely with the size (coherence length) of a periodic object. Longer cross-β fibrils will give rise to a sharper peak at ∼ 1.36 Å^-1^ (which arises from the 4.7 Å axial repeat of the cross-β structure). By subtracting the simulated broad tissue background (dashed curves in **Figure 1b**) from the data, the degree of tau fibrillation can be estimated from the magnitude of the residual intensity, highlighted in red in **Figure 1b**. Thus, scanning µXRD data enable us to map both the distribution of tau and the degree of tau fibrillation *in situ* across histological sections with micron-level resolution, making possible the correlation of structural information at the molecular level with cellular and tissue distributions.

### High resolution X-ray fluorescence imaging of light elements

Lighter elements such as S and P could not be accurately estimated at ID13 where XRF is excited at 13keV (far from their absorption edges) and XRF is collected in air (high reabsorption of low energy XRF signal). To determine the distribution of lighter elements, the same sample was reanalyzed in vacuum at beam line ID21 at ESRF.(26) For these experiments, the beam was focused to 0.3 μm (vertical) x 0.8 μm (horizontal) using Kirckpatrick Baez mirrors. Maps were acquired over regions of interest (ROI) of 99 µm x 99 μm, with steps of 1µm, and exposure time of 200ms. The X-ray beam was tuned to an energy of 4KeV to optimize signal from light elements including S, P and Ca. The use of a different sample holder for the ID13 and ID21 experiments precluded exact registration of the ID21 μXRF scans with the ID13 μXRD scans of the same sections. However, mapping of Ca distribution was carried out on both beam lines. This made it possible to use the Ca distribution collected at low X-ray energy as a guide to registration of scans taken at high X-ray energy simultaneously with the μXRD data.

## Acknowledgments

μXRD data were collected at ESRF ID13 and μXRF from ESRF ID21 under general proposal LS3086 and LS3300. This work was supported in part by the National Institute or Aging (R21-AG068972 and RF1AG079946 to LM) and the Rainwater Foundation (BTH). The Massachusetts Alzheimer’s Disease Research Center (MADRC) is supported by the National Institute on Aging (P30-AG-062421).

## Appendix: Molecular Polymorphism of tau aggregates in Pick’s disease

Jiliang Liu^a*^, Theresa Connors Stewart^b^, Bradley T. Hyman^b^, Manfred Burghammer^a^, Marine Cotte^a^, Lee Makowski^c^*.

a. European Synchrotron Radiation Facility, Grenoble 38043, France;

b. Department of Neurology, Massachusetts General Hospital, Charlestown, MA 02129, USA;

c. Bioengineering Department, Northeastern University, Boston, MA 02115, USA.

### The nanoscale packing of tau filaments determined by SAXS

The SAXS regime contains information on structural features of the scattering particles that have characteristic dimensions in the 1-100 nm scale. At this level, SAXS can be considered as scattering from a continuous distribution of electron density and approximated by a simplified model. Based on the structure of cross-β tau filament of Pick’s disease as determined by cryoEM (1), we constructed a parallelepiped model as shown in **Figure S4a**, with a cross section of 10 x 100 Å for the ‘narrow pick filament’ (NPF) and 10 x 200 Å for the ‘wide pick filament’ (WPF). The form factor of a parallelepiped can be calculated as:

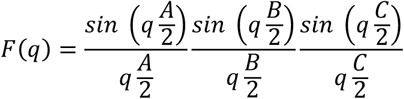

Considering the centroid symmetry of the parallelepiped, the 1D intensity profile can be calculated as the spherically averaged squared form factor as:

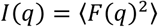

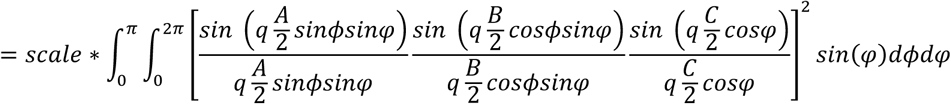

The slope of the SAXS intensity provides an approximation for the dimensions of these filaments. **Figure S4a)** and **b)** show a parallelepiped model with transverse dimension of 10 x 100 Å (A, B direction in **Figure S4a**) and 150 Å in longitudinal direction (C direction in **Figure S4a**) that gives rise to a profile with a slope comparable to the SAXS pattern from fibrillar tau (**Figure S4c**). This structural model is consistent with the NPF model of tau filament for Pick’s disease (Falcon et al. 2018). However, the features of the SAXS intensity pattern at 0.07 Å^-1^ and 0.2 Å^-1^ (**Figure 3**) indicate that, *in situ*, tau filaments further aggregate to form larger scale hierarchical structures within the lesion. The interference scattering arising from this higher order structure can be calculated as:

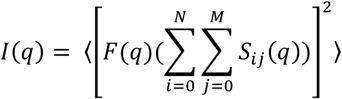

where 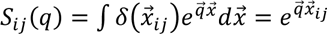 is the Fourier transform of the translation vector in real space. Thus, the spherically averaged intensity profile for the model of higher order aggregation of tau, shown in the Figure **S4c**, is calculated as:

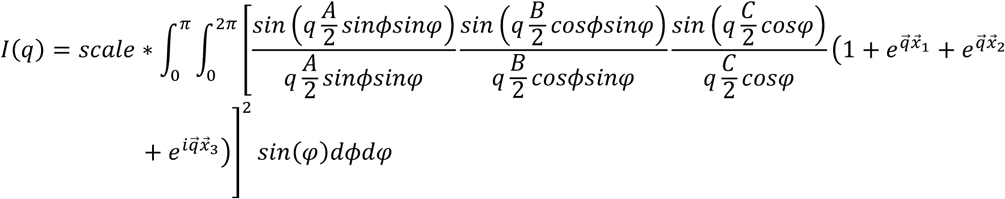

where x1= [30, 0, 0], x2 = [80, 0, 0,] and x3= [110, 0, 0]. **Figure S4d** shows that a SAXS profile calculated from a model including four parallelepiped fibrils with 10 x100 Å transverse structure and 20Å in the axial direction exhibits interference scattering consistent with experimental SAXS in both maximum position and log-log slope. However, the predicted sharpness of the calculated intensity is far greater than experimental data, suggesting polymorphism in the higher-order structural organization. The polymorphism in the the higher-order organization of tau fibrils was incorporated into the model as:

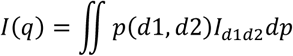

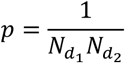

where d1 was given values of [ 20Å, 25Å, 30Å, 35Å, 40Å] and d2 [ 85Å, 92.5Å, 100Å, 107.5Å, 115Å]. The result of this integration is shown in **Figure S5**, which demonstrates that this results in a model for the hierarchical organization of tau in these NFTs that is consistent with observation. SAXS scattering includes contributions from other constituents of the lesions as well as that due to voids formed during dehydration of the tissue (17). In many cases, the contribution of other constituents of lesions to the observed scattering is closely similar to that observed in scattering from adjacent tissue. This ‘background’ scattering from tissue provides an estimate of the contribution of other constituents to that observed from lesions.

### The correlation between μXRD and μXRF

At high X-ray energy (13keV), the XRF spectrum of heavier elements was collected simultaneously with scanning μXRD. **Figure 1c** exhibits the averaged spectrum from three regions, one identified as tissue, one as low fibrillar tau and one as high fibrillar tau by the integral intensity of β-strands scattering at 1.36 Å^-1^ as detailed in **Figure 1b**. Qualitatively the density of sulfur (S), calcium (Ca), iron (Fe) and zinc (Zn) in the irradiated volume can be obtained by fitting the spectrum with elements determined from **Figure S6**.

**Figure S7** includes a superposition of the map of scattering of β-strands (from μXRD) with the maps of Zn and Ca. The map of scattering of β-strands has been normalized and transformed to an RGB image with μXRD as red channel and Zn as green channel, Ca as blue channel. In **Figure S7b** regions corresponding to the deposition of fibrillar tau has a white color, which indicates that the presence of high fibrillar tau with β-strands is associated with deposition of both Zn and Ca. In the Granular layer the strong correspondence of Zn and Ca with low fibrillar tau appears as cyan (light blue) instead of white.

### The high resolution XRF analysis

High resolution XRF image data was obtained by scanning the tissues with a 0.3 × 0.8 μm beam at ID21 ESRF using X-rays with an energy of 4 KeV to emphasize the signal from light elements such as phosphorous, sulfur and calcium. Data was collected on a square grid of 99 x 99 μm^2^ with step size of 1 μm^2^. A complete XRF spectrum such as that shown in **Figure S8** is collected at each scan position. The relative abundances of elements at each position are determined by fitting the XRF spectra with multi-Gaussian functions using PyMca software.

Due to differences in experimental setup, the registration of the high-resolution maps of calcium, phosphorous and sulfur in regions R1 to R4 vary slightly from that shown in **Figure S7**. However, the morphologies of cells, particularly those containing fibrillar tau, enabled the alignment of the high-resolution XRF maps with those in R1 to R4 in **Figure S7**. The high resolution XRF maps exhibit more details of the distribution of elements within the cell. Superposition of maps in **Figure S7** demonstrates that calcium and zinc content co-locate with both low-fibrillar and high-fibrillar tau. Region R4 of **Figure 5** and **Figure S7** shows that sulfur co-locates with fibrillar tau. Overlap of the maps of calcium, phosphorous and sulfur demonstrates that calcium and phosphorous are distributed throughout the pyramidal cells, which appears bright white in Region 3 and 4 in **Figure S9**.

---

**Figure S1.**
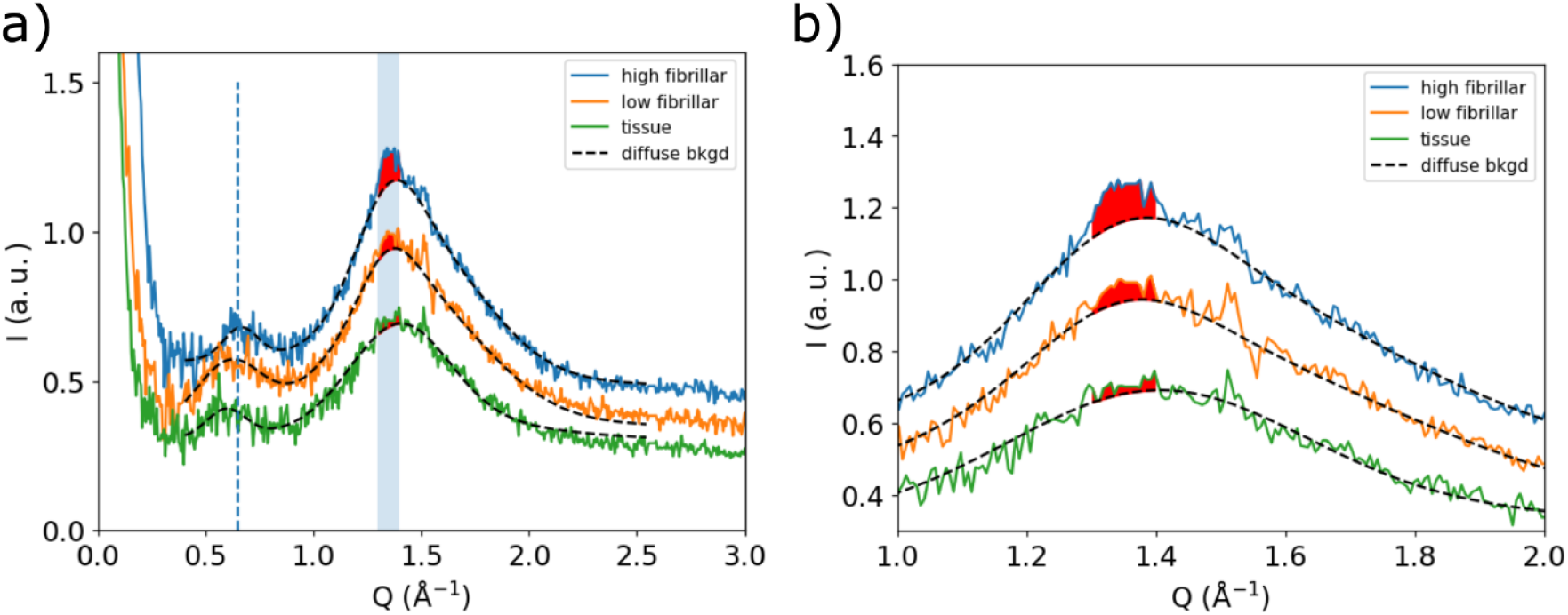
**a)** Comparison of diffraction from regions of a histological section containing only tissue (green), containing lesions dominated by low-fibrillar tau (orange) and containing lesions composed of fibrillar tau (blue). The smooth background curves (dashed lines) are calculated as a sum of Pseudo-Voigt functions and represent an estimate of the scattering of tissue which is comprised largely of fixed, partially denatured macromolecules. The vertical dashed line at Q = 0.65 Å^-1^ indicates the peak arising from structural features with characteristic dimension of ∼ 10 Å. The blue vertical bar highlights the Q range from 1.32 Å^-1^ to 1.4 Å^-1^, which corresponds to structural features with dimensions ∼ 4.7 Å. **b)** the deposition of tau enhances the intensity at ∼ 1.32 Å^-1^, with progressively greater levels of fibrillation giving rise to a sharper and more intense peak at that position. Scattering from low-fibrillar tau causes the broad, wide-angle peaks to shift from Q = 0.5 Å^-1^ to 0.65 Å^-1^ and Q = 1.42 Å^-1^ to 1.36 Å^-1^. The degree of fibrillation is estimated by subtraction of the diffuse background, resulting in a difference intensity (red) indicative of the degree of tau fibrillation. The broad scattering peak in the wide-angle regime spanning a region from 1.0 < Q < 2.0 Å^-1^ is generated by scattering from disordered, partially denatured and cross-linked macromolecules that make up much of the mass of the fixed tissue.

**Figure S2.**
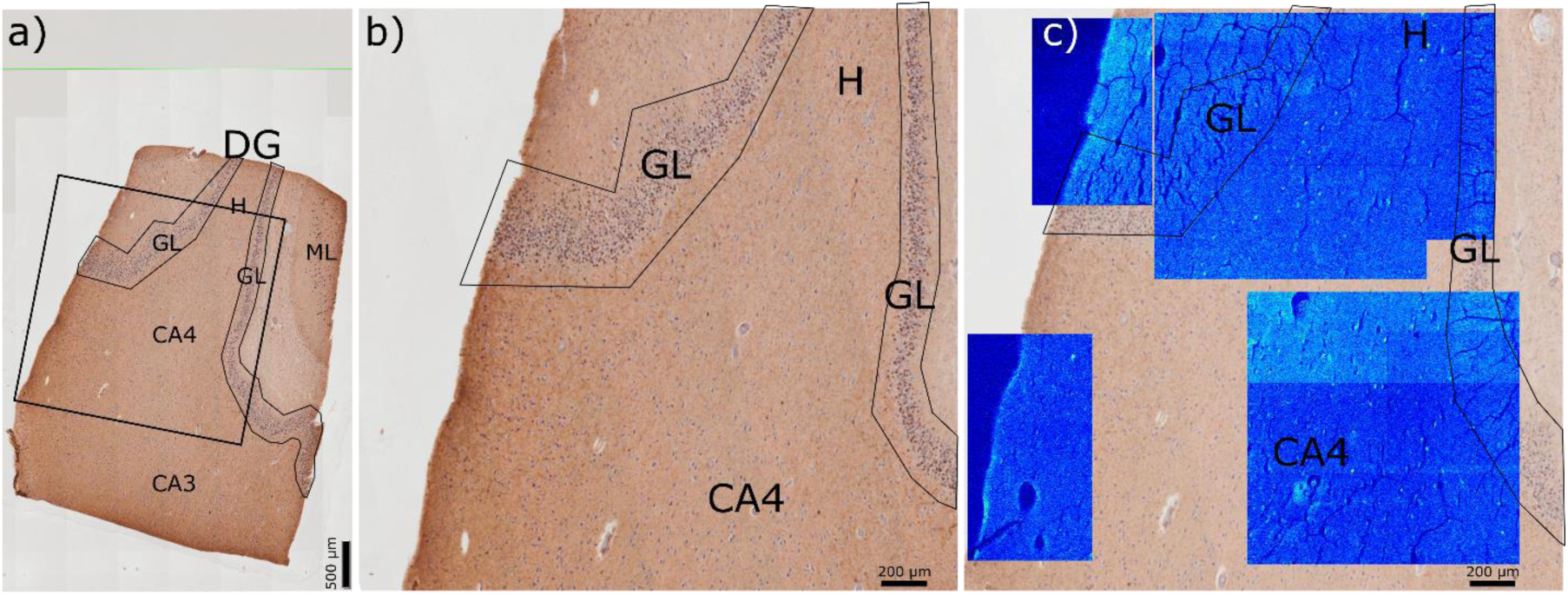
**a)** An immunostained serial section of tissue from Dentate Gyrus providing a larger anatomical context for the images in Figure 2 and exhibiting the distribution of tau and its association with tissue morphology. **b)** Enlargement of regions scanned by X-ray microdiffraction. **c)** Maps of the distribution of fibrillar tau determined by the intensity of the 4.7 Å peak (estimated as described in **Figure S1**) superimposed on the image of the immunostained serial section. Lighter blue indicates greater integral intensity. Additional analyses of these ROIs are detailed in Figure 2. Abbreviations in the figure: **GL**-Granular Layer, **H**-Hilus, **CA4**-Cornu Ammonis region 4.

**Figure S3.**
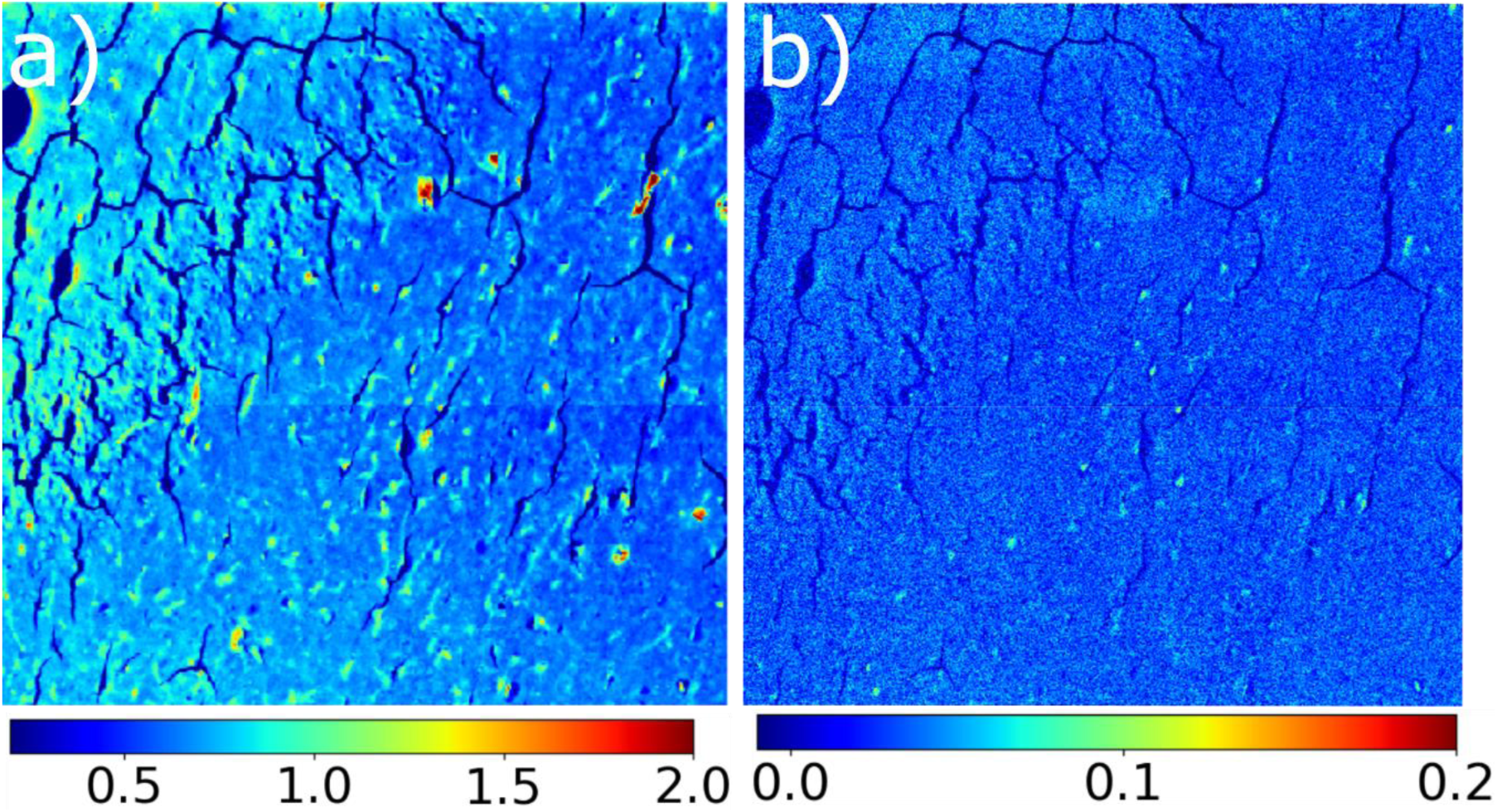
**a)** The map of the integral intensity of scattering from the Q range from 1.32 Å^-1^ to 1.4 Å^-1^ allows visualization of the distribution of molecular mass for the proteinaceous tissue in the Dentate Gyrus. The distribution includes many peaks that correspond to tau-containing lesions but does not distinguish between lesions with low or high levels of fibrillar tau. **b)** the distribution of fibrillar tau in the tissue as estimated by subtraction of diffuse tissue scattering as detailed in **Figure S1**. Lighter blue indicates greater concentration of fibrillar tau. Fibrillar tau is concentrated in punctate features prominent in (b) and is largely absent from the granular layer that occupies the upper right in these images.

**Figure S4.**
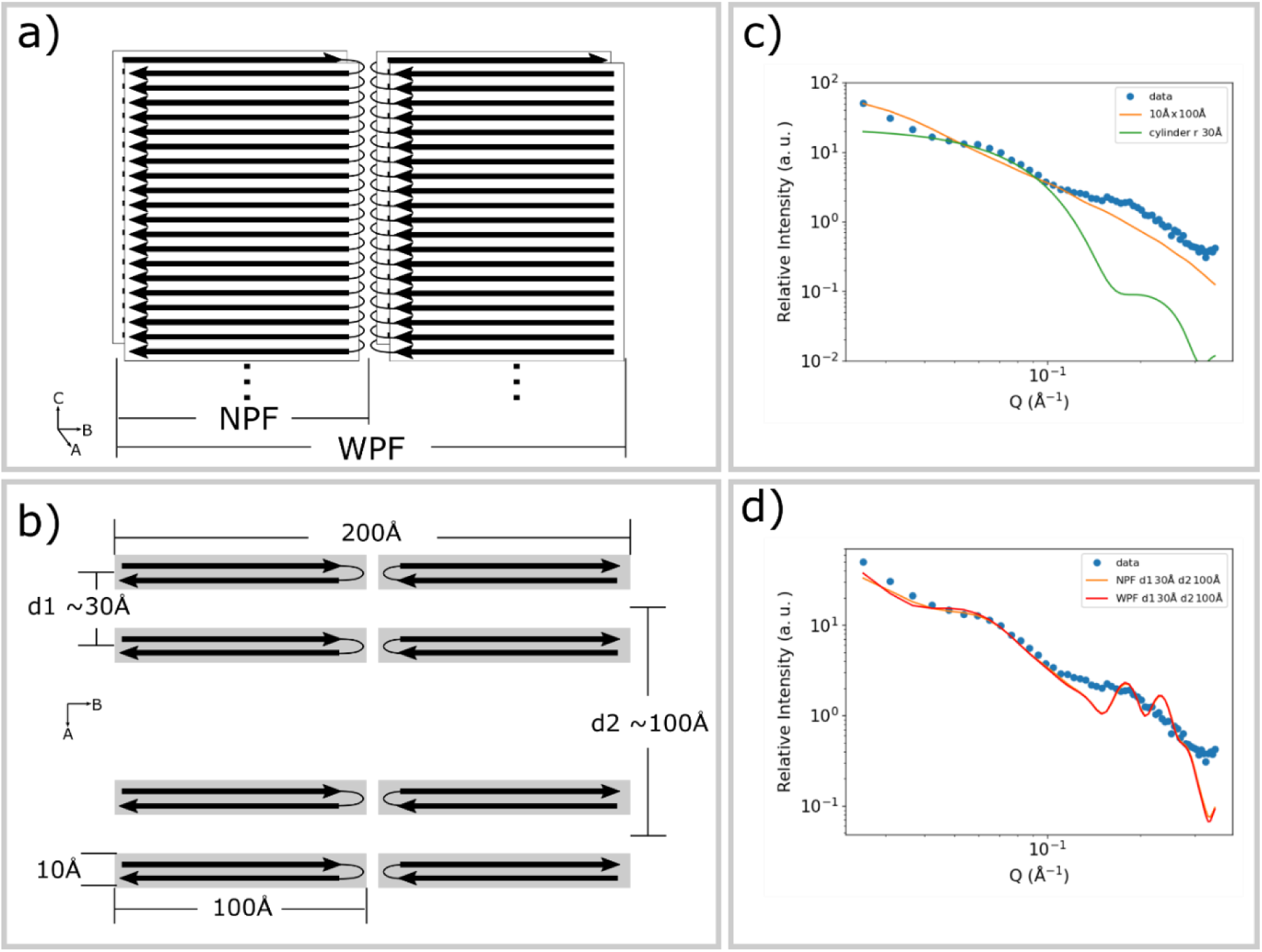
Constructing a hierarchical model of fibrillar Tau to fit the observed SAXS data. a) The fibril model is built on the basis of the structure of narrow fibrils (NF) determined by cryoEM. The wide fibrils are comprised of two narrow fibrils related to one another by a two-fold axis parallel to the fiber axis. b) Models constructed as hierarchical aggregates of narrow fibrils, including wide fibrils, exhibit organization on at least two length scales. The first is formed from the intra-fibril distance, d1, of ∼ 30 Å and the second from inter-fibril distances, d2, of ∼ 100 Å. c) SAXS calculated from the Narrow Fibril model (red) exhibits a slope comparable to that observed, but lacks the distinctive shoulders. A cylinder model (green curve) proposed earlier, (18), reproduces some of the SAXS features but does not predict the overall decrease of intensity as a function of Q. d) The calculated SAXS profile for these choices of d1 and d2 fits the first intensity shoulder well, but not the second.

**Figure S5.**
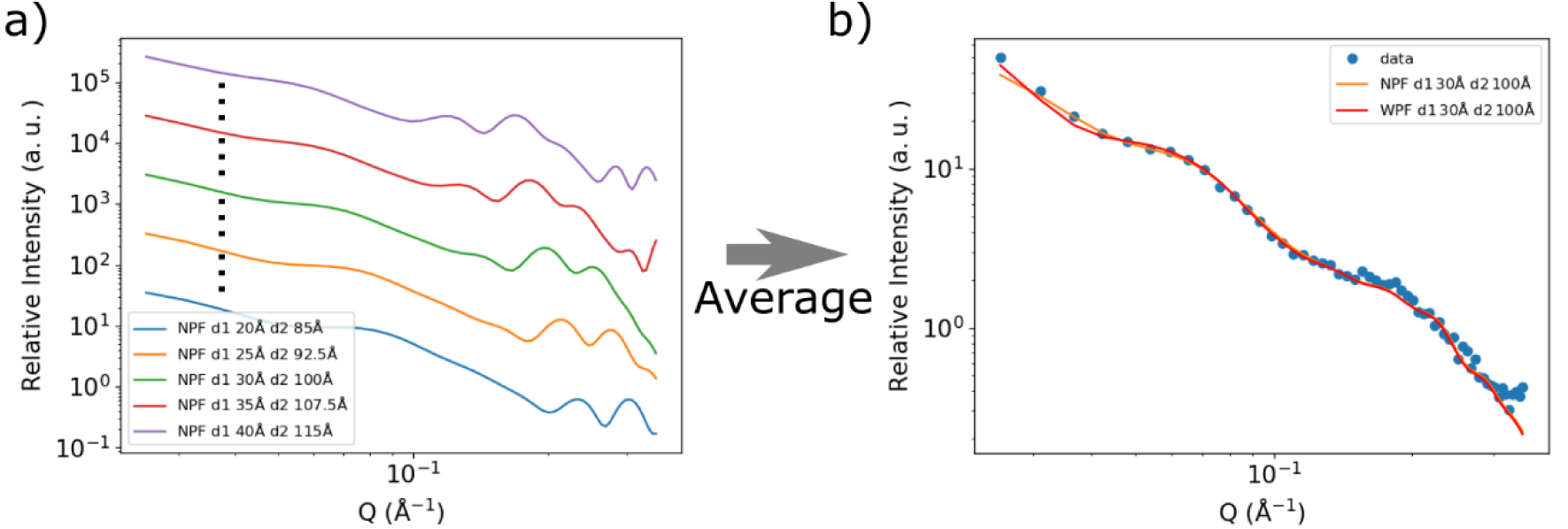
Adding polydispersity to the multi-scale hierarchical model for Tau filaments. a) For any given choice of d1 and d2, the calculated SAXS profiles include features not consistent with experiment. b) An ensemble of fibrillar aggregates constructed with d1 varying from 20 – 40 Å and d2 from 85 to 115 Å with equal probability, results in a SAXS pattern consistent with that observed *in situ* strongly suggesting that *in situ* the fibrillar aggregates are organized in a hierarchical structure that exhibits limited variation of fibril-fibril distances.

**Figure S6.**
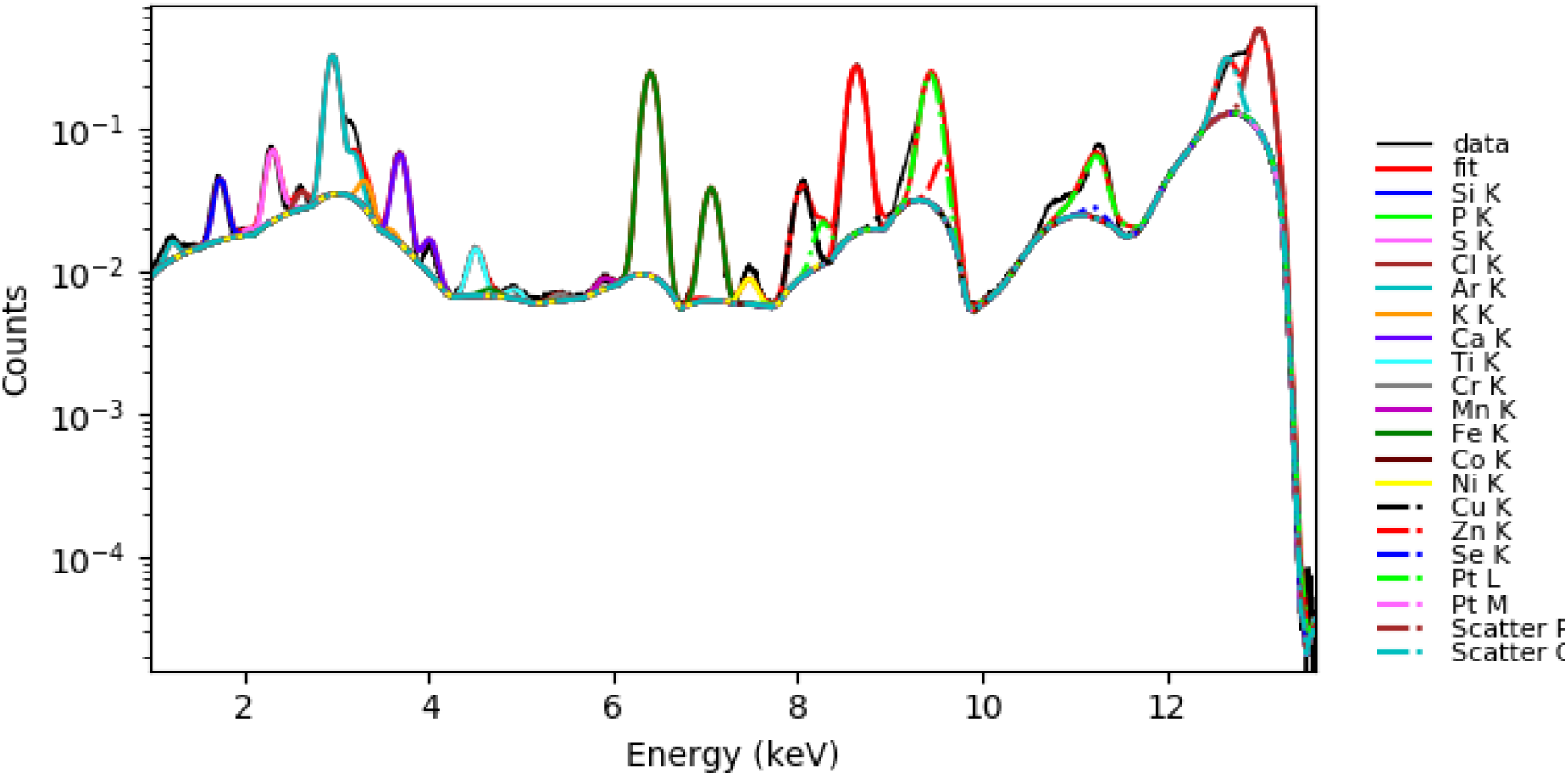
The averaged XRF spectrum from ID13 µXRF image map, with peak heights of individual elemental peaks estimated using PyMca.

**Figure S7.**
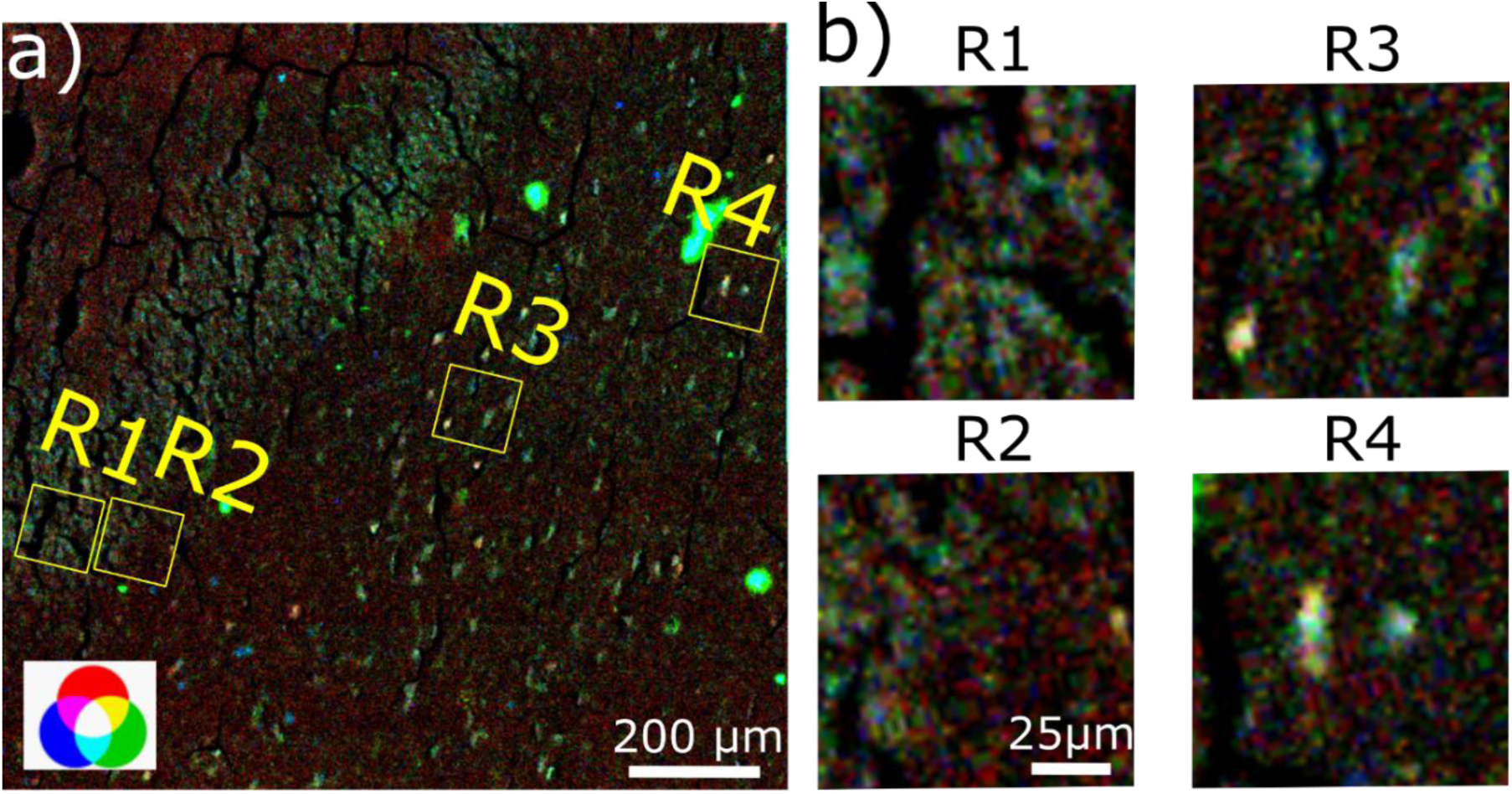
Superposition of the distribution of total tau estimated from the scattering at 4.7 Å with the distribution of Zn and Ca from XRF. (a) includes the entire ROI exhibited in Figure 2. (b) includes enlargements of four regions within that ROI. R1 and R2 in the granular layer exhibit strong co-deposition of Zn and Ca with low-fibrillar tau as visualized by the superposition of green and blue leading to cyan or light blue color in the enlargements on the right. In R3 and R4, in the hilus, places where fibrillar tau co-localizes with Zn and Ca appear as white. The label in **a)** shows the mixture of RGB colors.

**Figure S8.**
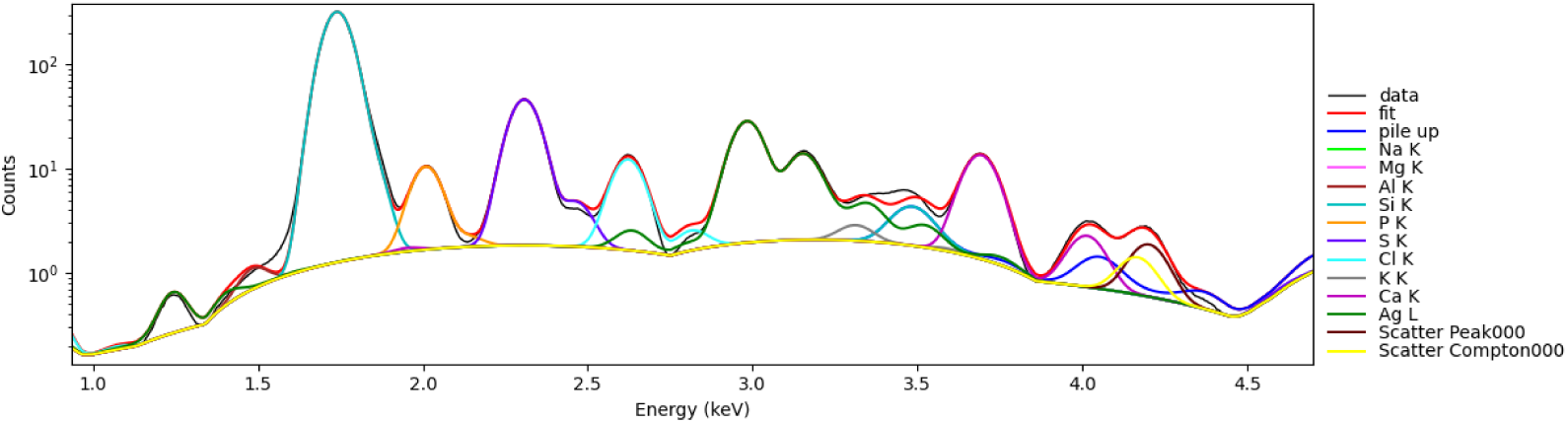
The XRF spectrum from ID21 µXRF image map, fitted using PyMca

**Figure S9.**
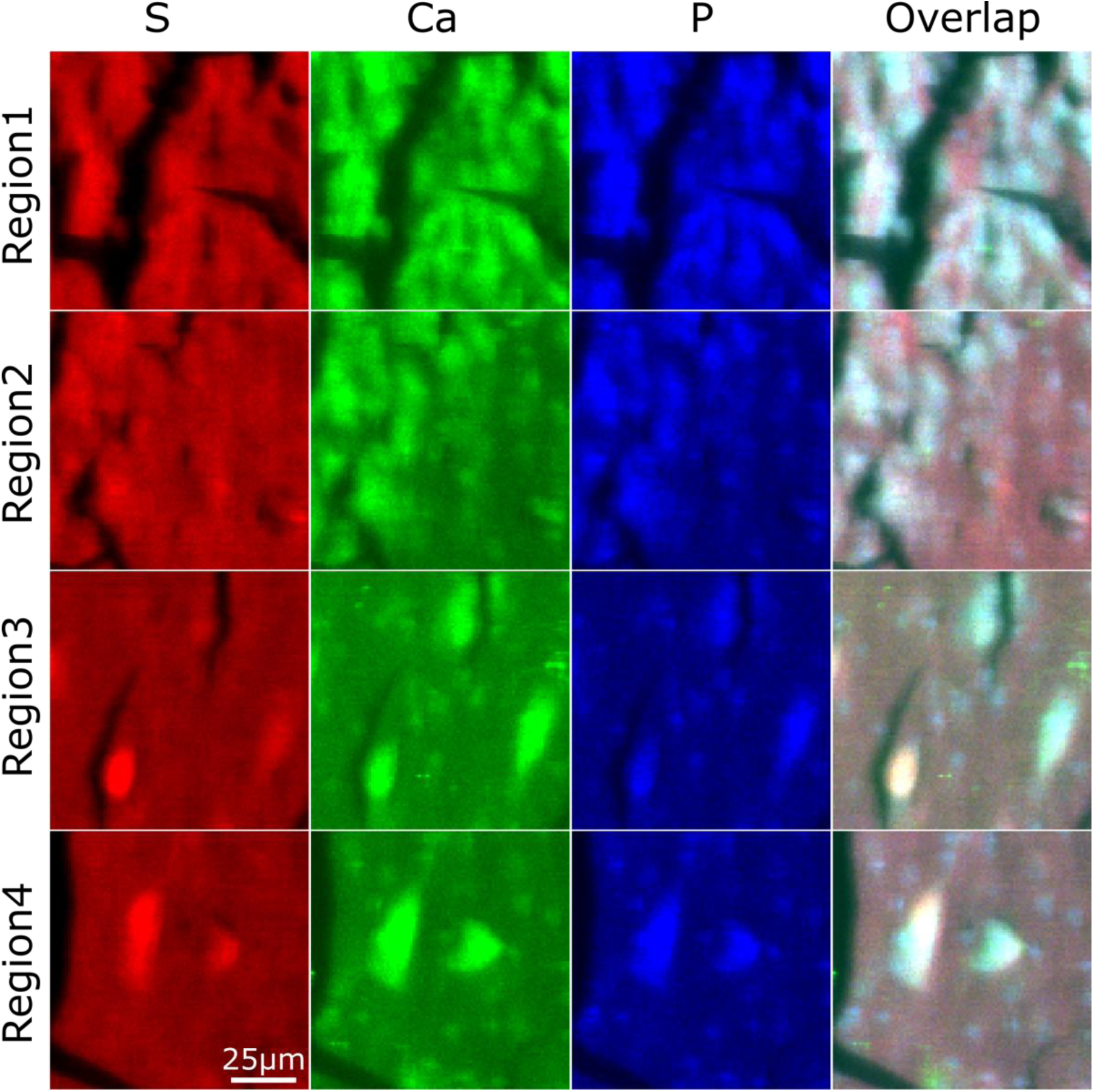
Comparison of the distribution of elements from high-resolution XRF data. Here, the maps in Figure 5 of main text are transformed to RGB format. The map of sulfur is transformed to the red channel, calcium to the green channel, and phosphorous to the blue channel. Both calcium and phosphorous exhibit deposition in all tau-containing lesions. Sulfur demonstrates strong correlation with calcium and phosphorous only in Regions 3 and 4, indicating a strong co-localization with fibrillar tau.

## References

1. Falcon, B., Zhang, W., Murzin, A. G., Murshudov, G., Garringer, H., Vidal, R.,…Goedert, M. (2018). Structures of filaments from Pick’s disease reveal a novel tau protein fold. Nature, 561(7721), 137–140.

2. Fitzpatrick, A. W., Falcon, B., He, S., Murzin, A. G., Murshudov, G., Garringer, H. J.,…Scheres, S. H. (2017). Cryo-EM structures of tau filaments from Alzheimer’s disease. Nature, 547(7662), 185–190.

3. Mate De Gerando, A., Welikovitch, L. A., Khasnavis, A., Commins, C., Glynn, C., Chun, J.,…Hyman, B. T. (2023). Tau seeding and spreading in vivo is supported by both AD-derived fibrillar and oligomeric tau. Acta Neuropathologica, 1-20.

4. Liu, J., Costantino, I., Venugopalan, N., Fischetti, R., Hyman, B., Frosch, M.,…Makowski, L. (2016). Amyloid structure exhibits polymorphism on multiple length scales in human brain tissue. Scientific reports, 6(1), 33079.

5. Berriman, J., Serpell, L. C., Oberg, K. A., Fink, A. L., Goedert, M., & Crowther, R. A. (2003). Tau filaments from human brain and from in vitro assembly of recombinant protein show cross-β structure. PNAS, 100(5), 9034–9038.

6. Kirschner, D., Abraham, C., & Selkoe, D. (1986). X-ray diffraction from inraneuronal paired helical fimanes and extraneuronal amyloid fibers in Alzheimer disease indcates cross-beta conformation. PNAS, 82(2), 503–507.

7. Carboni, E., Nicolas, J., Töpperwien, M., Stadelmann-Nessler, C., Lingor, P., & Salditt, T. (2017). Imaging of neuronal tissues by x-ray diffraction and x-ray fluorescence microscopy: evaluation of contrast and biomarkers for neurodegenerative diseases. Biomedical optics express, 8(10), pp.4331–4347.

8. Joppe, K., Nicolas, J., Grünewald, T., Eckermann, M., Salditt, T., & Lingor, P. (2020). Elemental quantification and analysis of structural abnormalities in neurons from Parkinson’s-diseased

9. Araki, K., Yagi, N., Ikemoto, Y., Yagi, H., Choong, C., Hayakawa, H.,…Nagai, Y. (2015). Synchrotron FTIR micro-spectroscopy for structural analysis of Lewy bodies in the brain of Parkinson’s disease patients. Sci. Rep., 5, pp.1–8.

10. Kawles, A., Minogue, G., Zouridakis, A., Keszycki, R., Gill, N., Nassif, C.,…Castellani, R. (2021). Differential vulnerability of the dentate gyrus to tauopathies in dementias. Acta neuropathologica communications, 11(1), 1.

11. Hyman, B., Van Hoesen, G. W., Wolozin, B. L., Davis, P., Kromer, L. J., & Damasio, A. R. (1988). Alz-50 antibody recognizes Alzheimer-related neuronal changes. Annals of neurology, 23(4), 371–379.

12. Amaral, D. (1978). A Golgi study of cell types in the hilar region of the hippocampus in the rat. Journal of Comparative Neurology, 182(5), 851–914.

13. Woelfle, S., & Boeckers, T. M. (2021). Layer-specific vesicular glutamate transporter 1 immunofluorescence levels delineate all layers of the human hippocampus including the Stratum lucidum. Frontiers in Cellular Neuroscience, 15, 789903.

14. Bashit, A. A., Nepal, P., Connors, T., Oakley, D., Hyman, B. T., Yang, L., & Makowski, L. (2022). Mapping the spatial distribution of fibrillar polymorphs in human brain tissue. Frontiers in Neuroscience, 16, 909542.

15. Wiśniewski, H. M., Coblentz, J. M., & Terry, R. D. (1972). Pick’s disease: a clinical and ultrastructural study. Archives of Neurology, 26(2), 97–108.

16. Liu, J., & Makowski, L. (2022). Scanning x-ray microdiffraction: In situ molecular imaging of tissue and materials. Current Opinion in Structural Biology, 75, 102421.

17. Nepal, P., Bashit, A. A., & Makowski, L. (2024). Characterization of sub-micrometre-sized voids in fixed human brain tissue using scanning X-ray microdiffraction. J. Appl. Cryst. DOI: 10.1107/S1600576724008987.

18. Schweers, O., Schonbrunn-Hanebeck, E., Marx, A., & Mandelkow, E. (1994). Structural Studies of Tau Protein and Alzheimer Paired Helical Filaments Show No Evidence for beta-Structure. The Jounral of Biological Chemistry, 269(39), 24290–24297.

19. Miller, L. M., Wang, Q., Telivala, T. P., Smith, R. J., Lanzirotti, A., & Miklossy, J. (2006). Synchrotron-based infrared and X-ray imaging shows focalized accumulation of Cu and Zn co-localized with β-amyloid deposits in Alzheimer’s disease. Jouranl of Structural Biology, 155(1), 30–37.

20. Goedert, M., Jakes, R., Spillantini, M., Hasegawa, M., Smith, M., & Crowther, R. (1996). Assembly of microtubule-associated protein tau into Alzheimer-like filaments induced by sulphated glycosaminoglycans. Nature, 383(60), 550–553.

21. Hasegawa, M., Crowther, R., Jakes, R., & Goedert, M. (1997). Alzheimer-like changes in microtubule-associated protein tau induced by sulfated glycosaminoglycans: inhibition of microtubule binding, stimulation of phosphorylation, and filament assembly depend on the degree of sulfation. Journal of biological chemistry, 272(52), 33118–33124.

22. Nishitsuji, K., & Uchimura, K. (2017). Sulfated glycosaminoglycans in protein aggregation diseases. Glycoconjugate Journal, 34, 453–466.

23. Lewkowicz, E., Jayaraman, S., & Gursky, O. (2021). Protein Amyloid Cofactors: Charged Side-Chain Arrays Meet Their Match? Trends in Biochemical Science, 46(8), 626–629.

24. Banani, S., Lee, H., Hyman, A., & Rosen, M. (2017). Biomolecular condensates: organizers of cellular biochemistry. Nature reviews molecular cell biology, 18(5), 285– 298.

25. Solé, V., Papillon, E., Cotte, M., Walter, P., & Susini, J. (2007). A multiplatform code for the analysis of energy-dispersive X-ray fluorescence spectra, Spectrochim. Acta Part B, 62, 63–68.

26. Cotte, M., Pouyet, E., Salome, M., Rivard, C., De Nolf, W., Castillo-Michel, H.,…Susini, J. (2017). The ID21 X-ray and infrared microscopy beamline at the ESRF: status and recent applications to artistic materials. Journal of Analytical Atomic Spectrometry, 32(3), 477–493.

## Appendix References

2. Nepal, P., Bashit, A. A., & Makowski, L. (2024). Small-angle X-ray microdiffraction from fixed human brain tissue exhibits power-law behavior that provides insights into the structural organization of neuropathological lesions. Biophysical Journal, 123(3), 135a.

3. Schweers, O., Schonbrunn-Hanebeck, E., Marx, A., & Mandelkow, E. (1994). Structural Studies of Tau Protein and Alzheimer Paired Helical Filaments Show No Evidence for beta-Structure. The Jounral of Biological Chemistry, 269(39), 24290–24297.

